# Relative model fit does not predict topological accuracy in single-gene protein phylogenetics

**DOI:** 10.1101/698860

**Authors:** Stephanie J. Spielman

## Abstract

It is regarded as best practice in phylogenetic reconstruction to perform relative model selection to determine an appropriate evolutionary model for the data. This procedure ranks a set of candidate models according to their goodness-of-fit to the data, commonly using an information theoretic criterion. Users then specify the best-ranking model for inference. While it is often assumed that better-fitting models translate to increase accuracy, recent studies have shown that the specific model employed may not substantially affect inferences. We examine whether there is a systematic relationship between relative model fit and topological inference accuracy in protein phylogenetics, using simulations and real sequences. Simulations employed site-heterogeneous mechanistic codon models that are distinct from protein-level phylogenetic inference models. This strategy allows us to investigate how protein models performs when they are mis-specified to the data, as will be the case for any real sequence analysis. We broadly find that phylogenies inferred across models with vastly different fits to the data produce highly consistent topologies. We additionally find that all models infer similar proportions of false positive splits, raising the possibility that all available models of protein evolution are similarly misspecified. Moreover, we find that the parameter-rich `GTR` model, whose amino-acid exchangeabilities are free parameters, performs similarly to models with fixed exchangeabilities, although the inference precision associated with `GTR` models was not examined. We conclude that, while relative model selection may not hinder phylogenetic analysis on protein data, it may not offer specific predictable improvements and is not a reliable proxy for accuracy.

## Introduction

When analyzing sequence data in evolutionarily-aware contexts, and in particular when inferring phylogenetic trees using modern statistical approaches, researchers must select an appropriate evolutionary model. The most common modeling framework for such applications follows a time-reversible continuous-time Markov process, usually considering either nucleotides, codons, or amino acids as states (Yang 2014; Arenas 2015). Since this framework’s introduction, a wide array of model parameterizations have been developed, ranging in complexity from the simple equal-rates Jukes-Cantor (`JC`) model (Jukes and Cantor 1969) where substitution rates among all states are equal, to the most complex form where all substitution rates are distinct (Tavare 1984), generally referred to as the `GTR` model. Additional levels of complexity beyond a model’s core substitution rates, such as incorporating among-site rate variation (ASRV), further increase the number of models from which practitioners can choose (Yang 2014).

To choose among dozens, if not hundreds, of available model formulations, the field has largely converged upon a strategy of relative model selection. For a given multiple sequence alignment, this approach systematically evaluates the statistical fit to the data for a set of candidate models using various metrics, most commonly information theoretic criteria (Posada and Buckley 2004). Such criteria include, for example, Akaike Information Criterion (AIC) and Bayesian Information Criterion (BIC), which provide a measure of goodness-of-fit to the data while penalizing models with excessive parameters that could lead to overfitting (Sullivan and Joyce 2005). Once available models are ranked by a given criterion, the model with the best fit to the data is subsequently specified during phylogenetic inference.

First popularized by the seminal software `MODELTEST` over 20 years ago (Posada and Crandall 1998), many different frameworks that perform relative model selection have been and continue to be developed (Darriba et al. 2011, 2012; Whelan et al. 2015; Kalyaanamoorthy et al. 2017; Darriba et al. 2019). Alongside this popularization has emerged a near-dogmatic mentality that employing the best-fitting model will increase the reliability, and potentially the accuracy, of inferences. Relative model selection has been described as “an essential stage in the pipeline of phylogenetic inference” (Arenas 2015) and is often viewed as a panacea to avoid model misspecification and biased inferences. While it is of course necessary to select a model of evolution for any analysis, whether relative model selection is the optimal procedure for doing so has been challenged in recent years (Luo et al. 2010; Brown 2014; Brown and Thomson 2018; Abadi et al. 2019). Even so, casual and potentially misleading remarks about the role of relative model selection abound across biological research fields. For example, the popular online database for HIV sequences, HIV LANL (https://www.hiv.lanl.gov/), contains an analysis option “Find-Model” to perform model selection on sequence data, leading with the header “Purpose: FindModel analyzes your alignment to see which phylogenetic model best describes your data; **this model can then be used to generate a better tree**” (emphasis added; https://www.hiv.lanl.gov/content/sequence/findmodel/findmodel.html). As most users of this feature are likely not experts in phylogenetic reconstruction, this phrasing will most likely interpreted to mean better-fitting models give better results by definition.

In spite of this pervasive attitude, there is no guarantee that the best-fitting model will infer the most accurate phylogenies. Indeed, relative model selection is inherently unable to determine whether a given model is reasonable to use in the first place. To circumvent this drawback, many have advocated for a shift in focus towards absolute model selection methods, or similarly tests of model adequacy (Bollback 2002; Brown 2014; Brown and Thomson 2018). Such approaches include analysis of posterior predictive distributions or hypothesis tests that ask whether the given inference model produces molecular properties (e.g. number of invariant sites, mean GC-content for nucleotide-level data, entropy, etc.) that mirror those seen in the data at hand (Goldman 1993a,b; Ripplinger and Sullivan 2010; Gelman et al. 2013; Brown 2014; Duchêne et al. 2015, 2016; Bollback 2002; Brown 2014; Duchêne et al. 2015; Höhna S. et al. 2017; Brown and Thomson 2018). Despite recommendations that model adequacy approaches should be used in conjunction with relative model selection, they have yet to see widespread adoption in large part due to their time-consuming, computationally-intensive nature (Duchêne et al. 2015; Brown and Thomson 2018). As such, the majority of researchers performing phylogenetic reconstruction will most often rely on relative model selection to justify using a given model.

Several lines of recent research have questioned the reliability of relative model selection in phylogenetic contexts. For example, Spielman and Wilke (2015b) showed that, in the context of identifying selection pressures from sequence data using codon models, AIC and BIC strongly prefer systematically biased models, and models that mitigate bias have relatively poorer fits to the data. Keane et al. (2006) observed that selecting protein models based on *ad-hoc* assumptions of biological relevance to the data may not result in improved inferences. Similarly, Spielman and Kosakovsky Pond (2018) found that the model employed when estimating site-level evolutionary rates in protein alignments has little, if any, effect on the inferred rates, with the primary exception that the simple equal-rates Jukes Cantor (JC) model has unique and previously unrecognized power to identify rapidly-evolving sites.

In the context of phylogenetic inference specifically, several studies have been conducted to investigate the consequences of employing different model selection criteria on nucleotide data. Ripplinger and Sullivan (2008) showed that, while different criteria choose different models, resulting phylogenies are not significantly different from one another, with most differences occurring at poorly-supported nodes. Most recently, Abadi et al. (2019) echoed and extended this insight to show that phylogenetic topologies inferred with the most complex time-reversible nucleotide-level model (`GTR+I+G`) did not differ significantly from the entirely uninformative `JC` model, even though `JC` is generally a poor fit to most datasets. In total, these studies have suggested that model selection itself may either be unnecessary or inadvertently lead to high confidence in biased results. A thorough examination of the practical ramifications of relative model selection is merited to reconcile these recent findings with the overarching sensibility that relative model selection is a fundamentally necessary component of phylogenetic analysis.

In this work, we explore whether there exists a systematic relationship between model fit and inference accuracy. In particular, we ask whether phylogenetic reconstruction performed with better-fitting models consistently leads more accurately inferred topologies compared to poorly-fitting models, specifically when conducting phylogenetic inference from protein data. Protein models are uniquely phenomenological compared to codon-level and nucleotide-level models because nucleotides, not amino acids, are the fundamental unit of evolution (Liberles et al. 2013; Jones et al. 2018). As such, precise evolutionary quantities such as mutation rate cannot be directly applied to protein data. From biological first principles, then, there is no mechanistic way to describe the evolutionary process when only protein data is available.

The simplest protein model, under the general time reversible framework, is described by a continuous-time Markov process with an instantaneous rate matrix, for the substitution amino acid *i* to *j, Q*_*ij*_ = *r*_*ij*_*π*_*j*_ scaled such that − ∑ *π*_*i*_*Q*_*ii*_ = 1. Parameters *r*_*ij*_ describe the substitution rate, or exchangeabilities, between amino acids *i* and *j*, and *π*_*j*_ represents the stationary frequency of target amino acid *j* (Yang 2014; Arenas 2015). These exchangeabilities represent the average propensity of each type of amino acid substitutions. As there are 189 such free parameters, assuming symmetric exchangeabilities, these values are rarely estimated from a given alignment itself. Instead, empirically-derived models with fixed exchangeabilities that have been *a priori* derived from hundreds or thousands of training datasets are most commonly applied. Early efforts to generate these models produced seminal matrices such as the Dayhoff model (Dayhoff et al. 1978), and statistical advances over time led to more robust models derived from substantially larger training datasets that were specifically intended for phylogenetic reconstruction. These include the commonly-used models `JTT` (Jones et al. 1992), `WAG` (Whelan and Goldman 2001), and `LG` (Le and Gascuel 2008), as well as certain specialist amino acid models like the chloroplast-sequence–derived `cpREV` model (Adachi et al. 2000) or Influenza-sequence–derived `FLU` model (Dang et al. 2010), for example. Unlike exchangeability parameters, the *π*_*j*_ parameters are more often estimated from the alignment at hand, either optimized during phylogenetic reconstruction or directly obtained by counting the amino acids in the alignment, known as the +*F* parameterization (Yang 2014).

While this modeling framework has become the default analysis choice for most users constructing trees from protein sequences, it ignores heterogeneous site-specific evolutionary constraints which are known to dominate protein evolution (Echave et al. 2016). It is possible to incorporate among site rate variation (ASRV), by scaling individual site rates according to a discrete Gamma distribution or similar (Yang 2014), but this procedure will still assume that the same evolutionary pattern governs each site in a given unpartitioned alignment. Other modeling approaches have been developed to more directly account for the pervasive heterogeneity in protein evolution, such as the Bayesian CAT model in `PhyloBayes` (Lartillot and Philippe 2004; Le et al. 2008) or mixture models which consider a distribution of individual matrices (Le et al. 2012; Arenas 2015). In spite of their known benefits, these models’ computational complexity and resource requirements have somewhat limited their adoption as the standard modeling framework standard in protein phylogenetics. For example, while the Bayesian CAT model is well-suited for long, multi-gene alignments, the underlying MCMC sampler generally cannot accommodate more than approximately 100 taxa, ultimately restricting the CAT model’s utility to phylogenomic analyses on a small number of sequences (http://megasun.bch.umontreal.ca/People/lartillot/www/phylobayes4.1.pdf). Therefore, in this study, we focus on the effects of applying more widely-used single matrix protein exchangeability models.

In addition, we may expect that relative model selection when applied to protein models has distinct behaviors versus when applied to nucleotide models. Relative model selection was first applied in phylogenetics with an eye towards identifying the model with the most suitable level of complexity for the data, in the context of nucleotide-level models (Posada and Crandall 1998). While nucleotide models often have different numbers of parameters, all empirical protein exchangeability models contain the exact same number of fixed exchangeability parameters, and options for increasing model complexity most entail adding very few additional parameters to account for phenomena such as ASRV or proportion of invariant sites. As such, the primary differences among empirical protein models emerge from different exchangeabilities and not from the complexity of substitution process itself. This study therefore also seeks to clarify the specific role that relative model selection plays in the context of protein sequence data, since many of the complexity concerns that exist for nucleotide models are not applicable.

Overall, we do not observe a strong, systematic relationship between relative model fit to the data and inference accuracy in resulting topologies. Except for the most relatively poorly-fitting models, protein models with drastically different fits to a given dataset infer highly consistent topologies. This study therefore demonstrates that relative model selection, as applied to protein data, may not have substantial effects on analysis so long as the most poorly-fitted models are not used. Therefore, we ultimately conclude that relative model selection is not an appropriate predictor of accuracy for this phylogenetic application.

## Methods and Materials

### Simulation approach

We adopted a simulation-based approach to assess whether employing models of different fit induce systematic shifts in the accuracy of inferred phylogenetic topologies. All simulations were conducted using the Python library `pyvolve` (Spielman and Wilke 2015a). We simulated alignments according to the site-wise codon-level mutation–selection (MutSel) model (Halpern and Bruno 1998). The instantaneous rate matrix for this model at a given codon site *k* is specified as

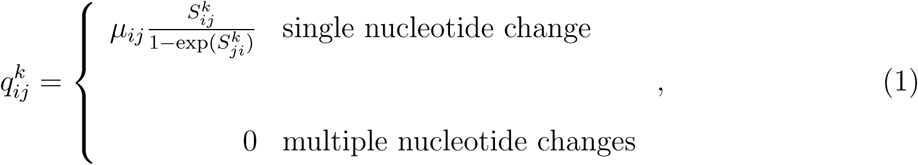

for a substitution from codon *i* to *j*, where *μ*_*ij*_ is the site-invariant nucleotide-level mutation rate, and 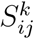 is the scaled selection coefficient for site *k*, which represents the fitness difference between codons *j* and *i* at site *k*, e.g. 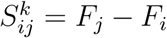 where *F*_*j*_ is the fitness of codon *j*. Each site *k* in a given alignment is therefore specified by a unique 61-length (the three stop codons are excluded) vector of codon fitness values.

We obtained site-specific codon fitness parameters used in simulations from four deep-mutational scanning (DMS) experiments (Table 1). DMS systematically determines the relative amino-acid preferences *P* for each site *k* in a real protein of length *L* (Firnberg et al. 2014). We converted these amino-acid preferences to fitness values as *F*_*a*_ = *log*(*P*_*a*_), where *F*_*a*_ is the fitness of amino acid *a*, and *P*_*a*_ is the experimentally-determined preference for amino acid *a* (Sella and Hirsh 2005). We assigned codon fitnesses based on these amino-acid fitnesses, assuming equal fitness among synonymous codons at each site *k*. We assumed site-invariant equal mutation rates among all nucleotides.

**Table 1:** Deep-mutational scanning experimental data used for MutSel simulations.

| Name | Description | Number of Sites | Reference |
| --- | --- | --- | --- |
| LAC | TEM-1 β-Lactamase | 262 | Firnberg et al. (2014); Bloom (2014b) |
| NP | Influenza H1N1 nucleoprotein | 497 | Bloom (2014a); Doud et al. (2015) |
| HA | Influenza H1N1 hemagglutinin | 564 | Thyagarajan and Bloom (2014) |
| HIV | HIV-1 Env Protein | 661 | Haddox et al. (2018) |

Each simulation was designed to mimic the evolutionary landscape of one of the real proteins given in Table 1. The length of each simulation therefore exactly matched the length of its respective originating protein. For example, all LAC-based simulations contained *L* = 262 codon sites, and the MutSel model operating at site *k* was parameterized by the experimentally-informed 61-length vector of codon fitnesses that corresponded to that position in the actual protein. This simulation strategies allows for highly realistic levels of ASRV as well as heterotachy (Spielman and Wilke 2015b; Jones et al. 2016, 2018). We simulated 20 alignment replicates for each set of DMS parameters along eight empirical phylogenies obtained from the literature (Table 2), resulting in a total of 640 simulated alignments. Because the MutSel model scales branch lengths to equal the expected number of neutral codon substitutions per unit time (Tamuri et al. 2012; Spielman and Wilke 2015a), all input phylogeny branch lengths were scaled up by a factor of three so that branch lengths would better approximate the number of amino-acid substitutions. Simulated codon-level alignments were translated to amino-acids before subsequent analyses.

**Table 2:** Empirical trees used for all simulations.

| Name | Number of taxa | Tree length* | Reference |
| --- | --- | --- | --- |
| Green plants | 360 | 24.67 | Ruhfel et al. (2014) |
| Ray-finned fish | 305 | 29.44 | Hughes et al. (2018) |
| Placental Mammals | 274 | 13.88 | dos Reis et al. (2012) |
| Aves | 200 | 5.21 | Prum et al. (2015) |
| Lassa virus | 179 | 6.62 | Andersen et al. (2015) |
| Spiralia | 103 | 25.01 | Marlétaz et al. (2019) |
| Opisthokonta | 70 | 20.92 | Ryan et al. (2013) |
| Yeast | 23 | 9.45 | Salichos and Rokas (2013) |

\*: Computed as the sum of branch lengths, measured in expected substitutions per site, from the phylogeny. In all simulations, these tree lengths correspond to amino-acid substitutions.

To complement these simulations, we performed a second set of simulations using the empirical protein model `WAG+I+G` (Whelan and Goldman 2001), where the proportion of invariant sites was set to 0.05. The discrete gamma distribution had 10 categories and a shape parameter of 0.8. The simulations represent a set of control simulations where each model used for inference have same mathematical general time-reversible form as does the generative simulation model. All control simulations were performed along the same phylogenies (Table 2), with 20 replicates for each of lengths 262, 497, 564, and 661 to act as analogs for MutSel simulations with LAC-, NP-, HA-, and HIV-derived parameters, respectively.

### Model selection and phylogenetic inference

For each simulated protein alignment, we employed `ModelFinder` in `IQ-TREE v1.6.8` (Nguyen et al. 2015; Kalyaanamoorthy et al. 2017) to determine the relative fit of standard protein exchangeability models using Bayesian Information Criteria (BIC). Because previous studies have shown that either most standard measures of fit when used in phylogenetics [BIC, Akaike Information Criteria (AIC), small-sample Akaike Information Criteria (AIC_*c*_), and decision theory (DT)] perform comparably in terms of impact on the emerging topology (Abadi et al. 2019), or that BIC may me somewhat more robust than other options (Luo et al. 2010), we focus on BIC alone in this study as the criterion for model selection.

We specified `ModelFinder` arguments that mimic behavior of the commonly-used `ProtTest` software (Darriba et al. 2011). `ModelFinder` examines a set of 21 commonly used protein models, as well as their respective parameterizations `+I` (proportion of uninformative sites), `+F` (using observed amino-acid frequencies), and `+G` (four-category discrete gamma-distributed ASRV), totalling 168 examined parameterizations. We ranked all evaluated models by their BIC score and identified five models ranging in goodness-of-fit for subsequent phylogenetic inference. Specifically, we identified the five models whose BIC scores most closely matched the five-number summary (minimum, first quartile, median, third quartile, and maximum) of the full distribution of BIC scores of all models evaluated for each alignment. We refer to the best-fitting model `m1`, the second-best fitting model `m2`, and so on for subsequent ranks along the five-number summary.

We then used `IQ-TREE v1.6.8` (Nguyen et al. 2015) to infer a phylogeny (e.g., optimize topology, branch lengths, and any additional free model parameters) using each of these five models, along with two additional models which were not considered in `ModelFinder`: The `JC` (Jukes and Cantor 1969) model, as well as the `GTR` model, where exchangeability parameters are optimized to the data during inference. Notably, these two modeling frameworks are generally unused in protein phylogenetics; the `JC` model is assumed to be overly simplistic and likely to underfit the data, and the `GTR` framework is presumed too parameter-rich and likely to overfit the data (Yang 2014). A full overview of the analysis pipeline for this study shown in Figure 1, using a single HA simulation replicate along the Placental Mammals tree as an example.

**Figure 1:**
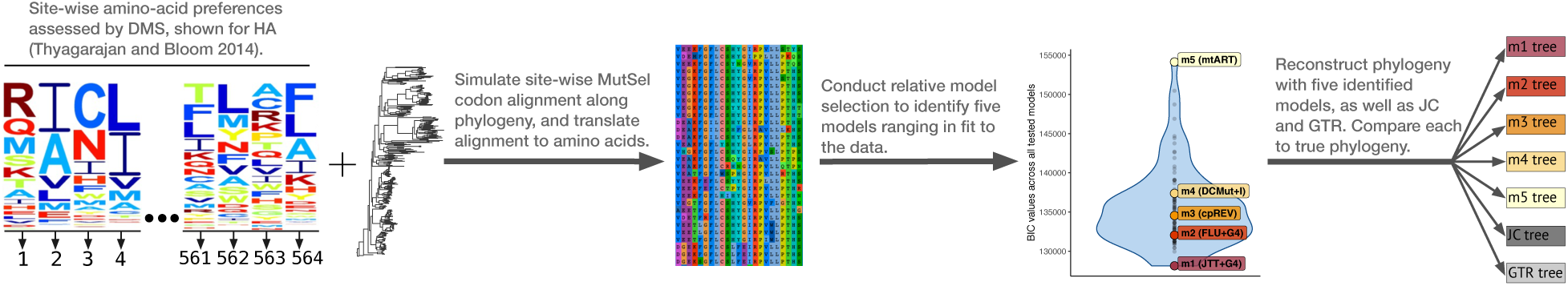
Flowchart describing simulation and phylogenetic reconstruction approach, for a single MutSel simulation replicate that uses HA site-wise preferences (Thyagarajan and Bloom 2014) along the Placental Mammals phylogeny as a representative example (dos Reis et al. 2012). Sequence logos depicting HA preferences were adapted from Thyagarajan and Bloom (2014). Model names shown in parentheses in the BIC distribution next to `m1-m5` correspond to the actual models used for this example HA simulation replicate. The violin plot represents the BIC scores for all 168 models evaluated by `ModelFinder` on this single alignment replicate, with highlighted model ranks indicated.

Calculation of Robinson-Foulds distance and other topological comparisons were performed using the Python library `dendropy v4.4.0` (Sukumaran and Holder 2010). Approximately unbiased (AU) topology tests (Shimodaira 2002) were performed in `IQ-TREE` by specifying the argument `-zb 10000 -au` to perform 10,000 RELL replicates (Kishino et al. 1990).

### Empirical data analysis

All empirical protein datasets were collected from the PANDIT database (Whelan 2006). 200 alignments (and corresponding PANDIT phylogenies) were randomly chosen from all PANDIT families where the “PANDIT-aa-restricted” set of sequences contained between 20– 500 (inclusive) sequences with between 100–1000 sites (inclusive). Relative model selection and phylogenetic inference were performed as described above.

### Statistical analysis and availability

All statistical analysis and visualization were performed in `R` (R Core Team 2017), making use of the `tidyverse` visualization and analysis framework (Wickham 2016; Wickham et al. 2019). Linear modeling was conducted using the `lme4` package (Bates et al. 2015), with corrections for multiple comparisons performed with `multcomp` (Hothorn et al. 2008). Significance throughout was assessed using a threshold of *α* = 0.01. All data and code, including results from all linear modeling analyses, are freely available from https://github.com/spielmanlab/aa_phylo_fit_topology and **will be deposited to Zenodo upon acceptance of the manuscript**.

## Results

### Model selection on simulated data

We began by simulating two broad sets of sequence alignments as described in *Materials and Methods*, which we term MutSel simulations and control simulations. Simulation is a powerful tool for studying the power and limitations of statistical methods, but the procedure is often highly confounded: A model must be employed to simulate data (generative model), but of course a model must also be chosen to analyze the data (inference model). If the inference model is accurately specified, we can expect strong inference model performance, particularly when using a consistent method such as maximum-likelihood estimation (Self and Liang 1987). However, when the generative and inference models correspond closely, it is easy to become overconfident about a model’s performance. Indeed, in real data analysis, any model used will be misspecified to a degree, some more than others. Therefore, to ensure that insights gained from simulations are not confounded by this logical gap, it is key to examine how models perform when the model is misspecified to the data. Such approaches have previously been shown, in evolutionary sequence analysis, to reveal unrecognized performance behaviors or biases in inference methods which would go unnoticed if the data met all model assumptions (Holder et al. 2008; Spielman and Wilke 2015b; Spielman et al. 2016; Spielman and Wilke 2016; Jones et al. 2016, 2018).

With both MutSel and control simulations, we can therefore ensure that results are not biased by similarities between generative and inference models. All control simulations use the `WAG` model which, like all empirical other protein models, can be decomposed into a vector of frequencies and symmetric matrix of exchangeabilities, but the MutSel model cannot be similarly decomposed. This is because, while both empirical protein models and MutSel satisfy detailed balance, they employ entirely distinct focal parameters: empirical models consider a site-invariant matrix of phenomenological exchangeabilities among amino-acids, but MutSel models, as employed here, consider site-wise codon fitness parameters coupled with site-invariant nucleotide-level mutation rates.

As `ModelFinder` does not evaluate the relative fit of the `JC` or the `GTR` models, we first examined their relative fit compared to `m1-m5` for each simulation (Figure S1). Across all MutSel simulations (Figure S1a), `JC` showed consistently poor fit and always ranked between models `m4` and `m5`. By contrast, the `GTR` model was consistently either the best-fitting model (for the larger NP, HA, and HIV simulations) or ranked in between the `m1` and `m2` models for most LAC simulations. Thus, `GTR` was a relatively high fit to the data for most MutSel simulations. For control simulations, the `GTR` model was always much lower-ranked, generally ranking between either models `m3` and `m4`, or between models `m4` and `m5` (Figure S1b). Thus, for control simulations, the parameter-richness of `GTR` was strongly penalized unlike for MutSel simulations. Similar to MutSel simulations, `JC` always ranked above the `m5` model, which was always the poorest-fitting model.

We generally expect that the best-fitting model determined by relative model selection (`m1`) should be the model that best reflects evolutionary properties of the given data. If this is indeed the case, we further expect that `m1` should be broadly consistent among simulations with the same set of DMS-derived parameters. Considering only the selected model matrix (i.e. both `LG+I` and `LG+F` would be considered the same model matrix, `LG`), we observed this expected trend in MutSel simulations (Figure 2). Interestingly, the `m1` model was mostly consistent for all simulations, regardless of which DMS parameter set was used, with either a `JTT`-based (Jones et al. 1992), `HIVb`-based (Nickle et al. 2007), or (for two LAC simulations) a `WAG`-based matrix emerging as the best-fitting model (Figure 2). Similarly, the model matrix `mtArt`, a model trained on arthropod-derived mitochondrial sequences (Abascal et al. 2006), was always the worst-fitting `m5` model across all alignments [with one exception of a single LAC simulation whose `m5` model was `mtMAM`, a model trained on mammalians mitochondrial sequences (Yang et al. 1998)] by a substantial BIC margin. Contrasting with the `m1` and `m5` models, a wide range of model matrices corresponded to `m2`, `m3`, and `m4` models, with between 3–13 model matrices observed at a given performance ranking (Table S1).

**Figure 2:**
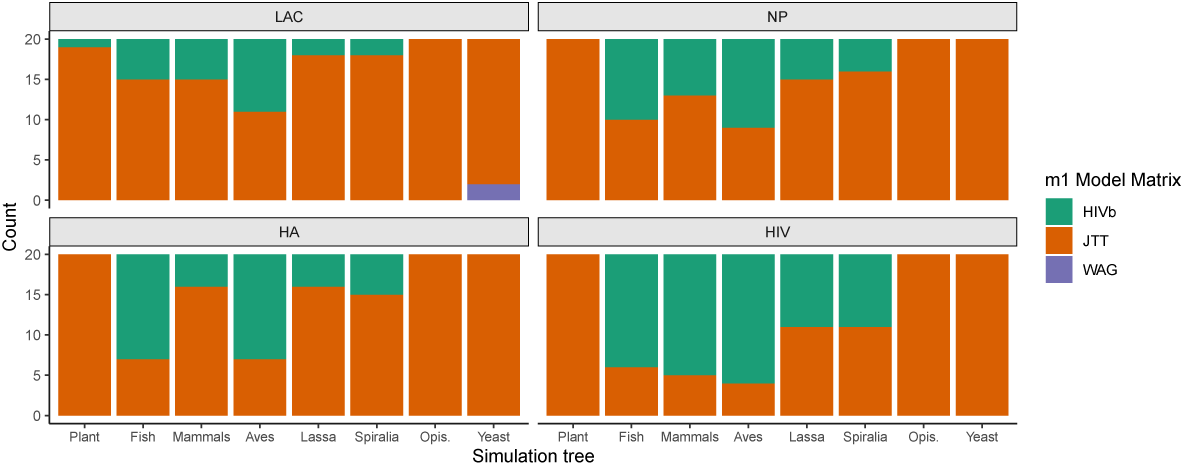
Best-fitting model (`m1`) matrix across MutSel simulations, where each column shows the selected model matrix for 20 simulation replicates. For visual clarity, the following abbreviations have been applied: Plant for Green plant, Fish for Ray-finned fish, Mammals for Placental mammals, Lassa for Lassa virus, and Opis. for Opisthokonta.

For all control simulations, a `WAG`-based matrix (always either `WAG+I+G` or `WAG+G`) always emerged as the `m1` model, matching the generative model and therefore representing a case of mostly accurate model specification. Similar to MutSel simulations, a wide variety of models corresponded to `m2`, `m3`, and `m4` models, with between 9–20 model matrices observed at a given performance ranking (Table S1). Further, the `m5` model for all control simulations was `mtMam` (Yang et al. 1998), again by substantial BIC margin.

There are several possible explanations for the overall similarity among `m1` model matrices for MutSel simulations. First, while these simulations accounted for realistic differences in protein-level selection, other simulation parameters could have induced overly-similar properties across alignments. For example, all simulations assumed symmetric and equal mutation rates among nucleotides, the same fitness among synonymous codons (no codon usage bias), and no indels (insertions/deletions). Alternatively, the similarity among `m1` model matrices may reflect inherent biases in experimentally-derived DMS preferences themselves. While DMS can recover local evolutionary constraints acting on each position in a protein, pooling all sites together may obscure protein-specific evolutionary signal, giving the appearance that entirely distinct proteins have more comparable evolutionary patterns (Ramsey et al. 2011). Indeed, it has been suggested that DMS-derived fitnesses may not always reflect true evolutionary constraints observed in nature due to the controlled laboratory conditions in which they are obtained (Haddox et al. 2016).

Finally, it is possible that `JTT` and `HIVb` happen to possess evolutionary information that generally represents protein evolution, in spite of the strong biological differences between the training datasets for each of these models. To probe relationship among these models further, we calculated the Pearson correlation between all pairs of model instantaneous rate matrices that `ModelFinder` considers (Figure S2). In fact, compared to all other models, `HIVb` model exchangeabilities are most strongly correlated with `JTT` exchangeabilities at *R* = 0.906. Furthermore, both `JTT` and `HIVb` models show the lowest correlations with `mtArt` (*R* = 0.792 and *R* = 0.645, respectively). The `HIVb` and `JTT` models are therefore much more similar than their origins suggest. That said, while `HIVb`’s strongest correlate is `JTT`, the `JTT` model itself is most strongly correlated with its variant `JTTDCMUT` (Kosiol and Goldman 2005) (*R* = 0.999), followed by models `VT` (Müller and Vingron 2000) (*R* = 0.954) (Müller and Vingron 2000), `WAG` (Whelan and Goldman 2001) (*R* = 0.935), `LG` (Le and Gascuel 2008) and `mtInv` (Le et al. 2017) (*R* = 0.916 and *R* = 0.914, respectively), and finally `HIVb` at *R* = 0.906. Even so, the strong correlation between `HIVb` and `JTT` may explain why these two specific models predominated as `m1` models.

### Inferred tree topologies show consistent distances from the true tree, regardless of relative model fit

To assess the relationship between model fit and phylogenetic topological accuracy, we first calculated the Robinson-Foulds (RF) distance between each inferred tree and the true simulation tree. To ensure consistent comparisons across simulation conditions, all RF distances were normalized by the maximum possible RF for the given phylogeny. Throughout, we use the acronym “nRF” to refer to normalized Robinson-Foulds distance.

If relative model fit is a reliable proxy for accurate protein phylogenetic inference, we should observe that nRF increases as model goodness-of-fit decreases, with the best-fitting model (`m1` or `GTR` for MutSel simulations, and `m1` for control simulations) inferring the most accurate tree topology. Our results from MutSel simulations, shown in Figure 3, did not convey this trend: nRF was remarkably consistent across inference models, with slight elevations apparent for `m5` models under certain simulation trees, most notably Opisthokonta. Instead, the most apparent trend we observed was that nRF decreased as the number of sites in the dataset increased, with HIV simulations showing much smaller nRF values compared to HA, NP, and LAC simulations. Results from control simulations (Figure S3) broadly echo these trends, which indeed suggests that sequence length is a primary driver for decreased nRF. Moreover, the overarching similarity between MutSel and control (whose simulations employed `WAG+I+G` with 10 rate categories) demonstrates that these results hold under both circumstances of strong model misspecification (MutSel results) and circumstances of little-to-no model misspecification (control simulations, namely `m1` results).

**Figure 3:**
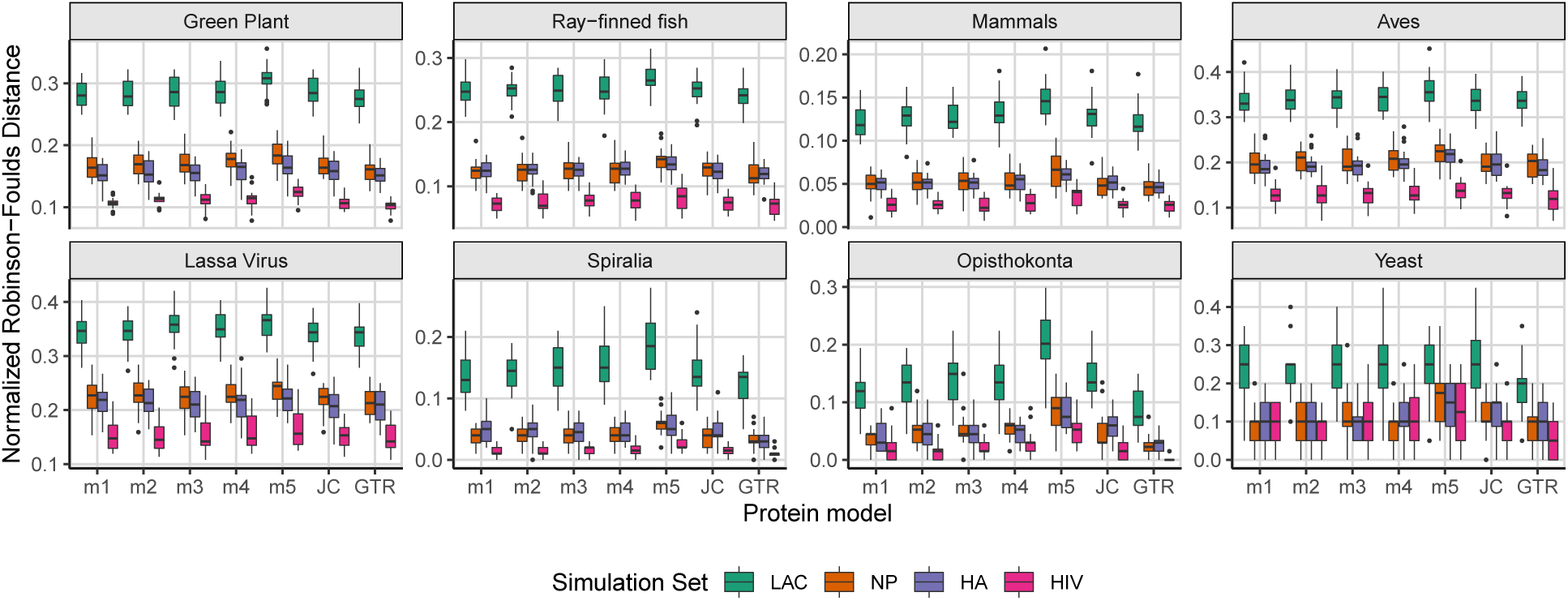
Normalized Robinson-Foulds distance (nRF) between tree inferences and the respective true tree from all MutSel simulations. Each boxplot represents the distribution of nRF values for 20 simulation replicates.

We fit a mixed effects linear model to determine the specific influence of protein model fit on nRF in MutSel simulations, specifying nRF as the response, the protein model (`m1`-`m5`, `JC`, and `GTR`) as a fixed effect, and the simulation tree and DMS parameterization each as random effects. We performed a Tukey test to evaluate pairwise differences in mean nRF among protein models. Trees inferred with models `m1`, `m2`, `m3`, `m4`, and `JC` (with the single exception of the `m1-m4` comparison) did not have significantly different nRF values, suggesting highly comparable performance among most models regardless of fit. Trees inferred with `m4` models did show a significantly larger nRF compared to `m1` trees, but with a vanishingly-small effect size of merely 0.7%. Trees inferred with `GTR` showed significantly smaller nRF compared to all other models, and trees inferred with `m5` models showed significantly larger nRF compared to all other models. All significant differences detected had exceedingly small effect sizes, with the largest difference in nRF of 3.3% from the comparison between `GTR` and `m5`. As such, the average nRF improvement from applying the best-fitting model compared to the worst-fitting model is, at most, roughly 3%, ultimately demonstrating that relative model fit does not have a strong systematic influence on phylogenetic inference accuracy. Analogous linear models performed on the control simulations again were largely consistent, with only extremely small effect sizes (at most 2.2%) for all models with significantly different nRF.

### All models infer similar amounts of strongly-supported but incorrect splits

Although model fit did not substantially affect nRF in any simulations, very few inferences exactly matched the true tree (RF distance of 0). Out of the total 4480 tree inferences conducted for each simulation set (seven trees inferred for each of 640 simulated alignments, for Mutsel and control each), only 130 Mutsel inferences and 412 control inferences achieved RF distance of 0 (Table S2). All of these inferences, for both MutSel and control simulations, were from simulations along either the Spiralia, Opthisthokonta, or Yeast phylogenies, the three trees with the fewest number of taxa (Table 2). For MutSel simulations, only NP (23/130), HA (20/130), and HIV (87/130) alignments achieved phylogenies with nRF of 0. For control simulations, all simulation lengths achieved phylogenies with nRF of 0 (LAC: 28/417, NP: 90/417, HA: 132/417, and HIV 162/417). Notably, all models, including `m5`, were able to reach the true tree for at least one replicate, for both MutSel and control simulations, although less commonly than other models. Further, the `GTR` model most frequently yielded the true true (44/130) for all MutSel simulations, but only yielded the true tree for 57/417 control simulation inferences. The `m1` model most frequently yielded the true tree (88/417) for control simulations.

However, RF distance is a notoriously conservative metric that considers only presence or absence of nodes without considering their uncertainty, i.e. the level of support for inferred nodes under a given inference model. If differing splits are poorly supported, RF will over-state the distance between trees being evaluated. By contrast, differing splits with strong support represent more problematic deviations from the true tree.

We therefore evaluated bootstrap support for each inferred phylogeny using the ultra-fast bootstrap approximation (`UFBoot2`) implemented in `IQ-TREE` (Minh et al. 2013; Hoang et al. 2017). Because `UFBoot2` is presumed a less biased measure compared to the standard nonparametric bootstrap, it necessitates a somewhat different interpretation such that nodes with ≥ 95% support are considered highly reliable (Minh et al. 2013; Hoang et al. 2017). In addition, this threshold of 95% for identifying supported splits is expected to correspond to a false positive rate (FPR) of 5%.

For each inferred tree, across all models, we evaluated whether each inferred node was accurate (present in the true tree) and whether each inferred node was strongly supported (`UFBoot2` ≥ 0.95) under the given model. We evaluated the FPR as well as the accuracy at this `UFBoot2` threshold. For these calculations, we specifically considered a given node as “true” if it was present in the true tree, and we considered a given node as “false” if it was not present in the true tree. We considered a given node as “positive” if its `UFBoot2`≥ 0.95, and we considered a given node as “negative” if its `UFBoot2`< 0.95. FPR is calculated as 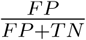, where *FP* is the number of false positive nodes and *TN* is the number of true negative nodes. The accuracy is calculated as 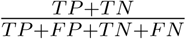, where *TP* is the number of true positive nodes, *TN* is the number of true negative nodes, *FP* is the number of false positive nodes, and *FN* is the number of false negative nodes. The resulting classification metrics, specifically for HA MutSel simulations, are shown in Figure 4. Corresponding results for LAC, NP, and HIV MutSel simulations are in Figures S4 and S5, and analogous results for all control simulations are shown in Figures S6 and S7. Trends in results from control simulations were again largely consistent with those from MutSel simulations.

**Figure 4:**
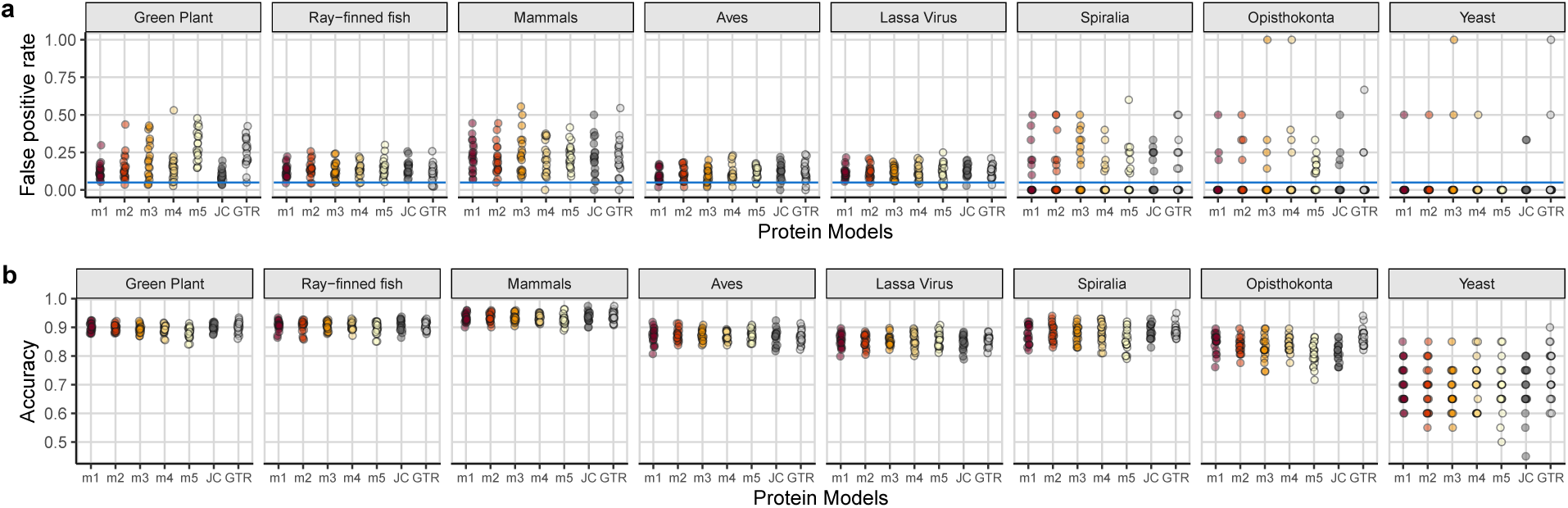
a) False positive rate (FPR) in inferred splits, for HA MutSel simulations, using 95% `UFBoot2` as a threshold. Each point represents the FPR of a single simulated alignment. The horizontal line in each panel is the *y* = 0.05 line, representing the expected FPR. b) Proportion of accurately-classified splits in tree inferences, for HA simulations, using 95% `UFBoot2` as a threshold. Each point represents the accuracy of a single simulated alignment.

Results for this analysis agreed with those from nRF analysis: Protein models ranging in fit to the data yielded similar levels of support, with both FPR and accuracy being remarkably similar across all inference models. However, the FPR was not well-bounded at the expected value of 5%, suggesting that `UFBoot2` may not be as robust to model violations as has been previously presumed (Hoang et al. 2017). That said, FPR was additionally not well-bounded for control simulations, even those whose trees were inferred with the correctly-specified WAG model (Figure S6). Further investigation of the sensitivity of this approximate bootstrap measure to model misspecification, and more generally, may therefore be merited. It is worth noting, however, that the overall percentage of false positive splits out of all splits 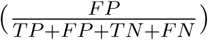, rather than FPR, was generally below 5% for all simulations, meaning that while FPR is somewhat high, the inferred trees are certainly not dominated by false-positive nodes (Figure S8 for MutSel simulations, and Figure S9 for control simulations).

We again analyzed this data with two linear models, considering either FPR or accuracy as the response, both with protein model as a fixed effect and DMS parameterization and tree as random effects, using a Tukey test to perform pairwise comparisons across protein models. Separate models were conducted for MutSel and control simulations. Similar to results from nRF analyses, MutSel modeling results showed no significant difference in FPR among trees inferred with all models `m1`, `m2`, `m3`, `m4`, and `JC`. By contrast, the `GTR` model had significantly larger FPR compared to `m1`, `m2`, and `JC`, and the `m5` model had significantly larger FPR compared to `m1`, `m2`, `m3`, and `m4`. There was no significant difference in FPR between `m5` and `GTR`. Even so, the largest effect size for any comparison was still extremely small at a maximum of 3.7% (for the comparison between `m1` and `m5`). Accuracy was not significantly different among models `m1`, `m2`, `m3`, `m4`, and `JC`, but `GTR` did show significantly higher accuracy compared to all other models, and `m5` showed significantly lower accuracy compared to all other models. Again, however, effect sizes for all significant comparisons were very modest, with at most a 2.3% difference in accuracy for the comparison between `GTR` and `m5` models, and analogous linear modeling results for control simulations were generally consistent.

In total, consistent with nRF analysis, protein models ranging in fit to the data performed highly comparably, with the worst-fitting `m5` model only producing marginally worse results than other models. That neither FPR nor accuracy were significantly different among `m1`, `m2`, `m3`, `m4`, and `JC` models provides further evidence that relative model fit does not have substantial bearing on phylogenetic inference from protein data. These results were also robust to the particular simulation strategy.

### Most inferred trees fall in the confidence set of trees under the m1 model

We next performed a series of AU (approximately unbiased) tests of tree topology to assess whether the observed topological differences represented significant deviations from the `m1` phylogeny (Shimodaira 2002). For each alignment, we performed an AU test to compare the alignment’s eight associated topologies: seven inferred trees and the true tree.

Across all MutSel simulations, we identified exceedingly few instances where any inferred phylogeny fell outside the `m1` confidence set, at a threshold of *P* < 0.01 (Table S3). All trees inferred with models `m2`, `m3`, `m4`, and `JC` fell inside the respective `m1` confidence set of trees (all *P* ≥ 0.044). By contrast, for 19 simulations (4% of total), the `m5` tree uniquely fell outside the `m1` confidence set, and for a single simulation replicate the `GTR` tree uniquely fell outside the `m1` confidence set. Most importantly, the true tree was in the `m1` confidence set for all but 31 simulations (6.5% of total), most commonly HA simulations. That true trees most commonly fell in the `m1` confidence set was somewhat surprising, given the substantial topology differences between inferred and true topologies (Figure 3). It is therefore a distinct possibility that trees with substantial topological differences may have more similar than anticipated likelihoods, an issue which merits future investigations. For control simulations, only 22 inferences fell outside the `m1` confidence set of trees: 17 inferred with `m5` and 5 inferred with JC. Only 12 true trees fell outside the `m1` confidence set.

### Analysis of natural sequence data reveals similar levels of consistency across models

We next examined how relative model fit affects phylogenetic inference for a set of natural protein sequence alignments. Because such analyses cannot truly assess inference accuracy (as the true phylogeny is unknown), we asked whether protein models ranging in goodness-of-fit to the data yielded consistent or significantly different topologies. Furthermore, although the `JC` and `GTR` models performed well on simulated data, it is possible that these results were an artifact of the relative simplicity of simulated data compared to the complexity of natural sequence data. Examining how these protein models perform on real data is therefore crucial to properly contextualize their strong performance in simulations.

We randomly selected 200 protein alignments from the PANDIT database, considering only those with 20–500 (inclusive) sequences and 100–1000 sites (inclusive). As with simulated data, we determined the five protein models which most closely matched the BIC quartiles, and we used each model, as well as `JC` and `GTR`, to infer a phylogeny. The exact protein models identified at the BIC quartiles were substantially more varied compared to selected models for simulated data (Table S4), although over 75% of datasets selected an `LG`-based `m1` model (Figure 5a). This strong bias towards LG likely reflects that the `LG` model itself was trained using PFAM alignments, the source for the PANDIT database (Whelan 2006; Le and Gascuel 2008). Similar to results from simulated data, the vast majority of `m5` models were either mtArt (Abascal et al. 2006) or mtMam (Yang et al. 1998). The general poor fit of certain mitochondrial models for both simulated and natural sequence data here may reflect the highly unique nature of the data on which these models were originally trained.

**Figure 5:**
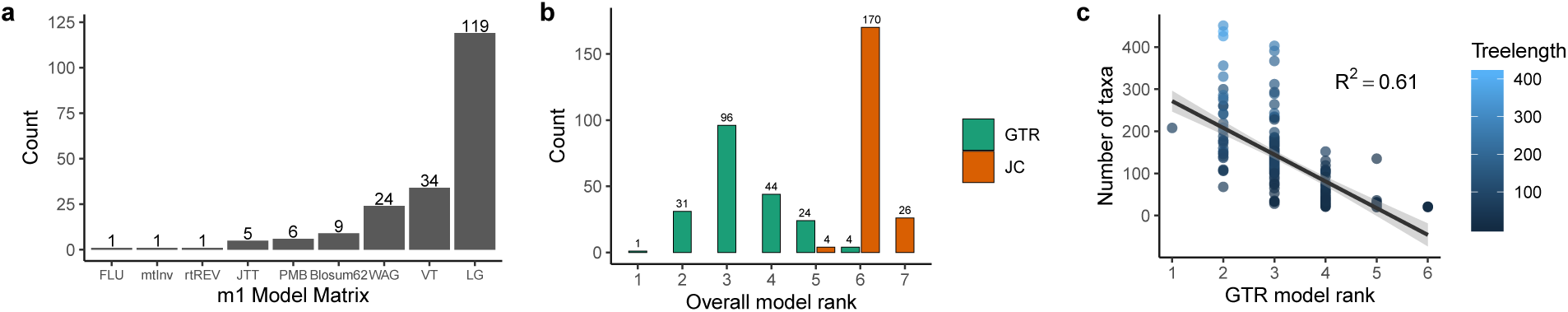
Model selection results on 200 PANDIT alignments. a) Best-fitting model (`m1`) matrix across PANDIT alignments. b) Relative rank, among all seven inference models, for the `JC` and `GTR` models, which were not considered by `ModelFinder`. c) Scatterplot showing key features in PANDIT datasets which predicted the relative `GTR` model rank within the seven inference models employed. Each point represents a PANDIT alignment, and treelength represents the sum of branch lengths each alignment’s respective `GTR`-inferred phylogeny.

Starkly contrasting with simulated MutSel alignments, but similar to the control simulations, the `GTR` model only emerged as the best-fitting model for a single PANDIT alignment (Figure 5b). Even so, `GTR` was generally a better fit to each dataset than were `JC` and `m5` models. As the PANDIT alignments analyzed here were substantially more sparse compared to simulated alignments, with percent of gaps and ambiguous amino acids ranging from 19%–79% across alignments, the relatively poorer fit of the `GTR` model is not unreasonable. We tested whether certain features of the alignments, including percent of ambiguous characters, number of taxa, length of alignment, and/or treelength (sum of inferred branch lengths) could explain the relative rank of the `GTR` model among the seven models examined for each alignment. Step-wise linear model selection using *R*^2^ showed that the best model to explain `GTR` rank was `GTR_rank` ∼ `number of taxa*treelength`, with *R*^2^ = 0.61 (Figure 5c). Thus, `GTR` tended to be a better relative fit for more informative alignments.

We examined to what extent tree topologies inferred across protein models were consistent with one another using two separate analyses: i) an all-to-all comparison of Robinson-Foulds distances, and ii) AU tests for each inferred set of trees to assess whether they fell in the respective `m1`-inferred tree’s confidence set. In Figure 6a, we show the distributions of nRF distances across each pair of models, with the median value shown for each distribution. Overall, the mean nRF between `m1` and `m2` trees was significantly lower than all other comparison distributions (*P* < 0.01), but we emphasize that the `m1`-`GTR` comparison showed the second lowest median nRF difference. There was virtually no other difference in average nRF for most comparisons among `m1`, `m2`, `m3`, `m4`, and `GTR` models. As such, while there are clear topological differences among trees inferred across these models, no single protein model of these five stood out as yielding substantially different topologies. These results are highly consistent with those from simulations and again suggest that relative model fit does not systematically affect inferred tree topologies.

**Figure 6:**
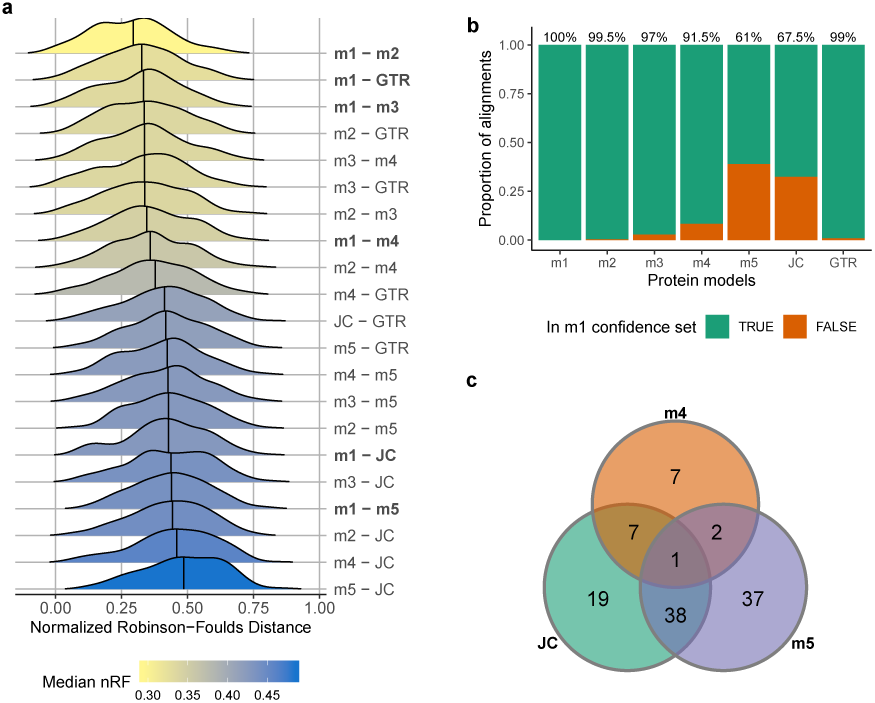
Results from PANDIT analysis. a) Distributions of nRF differences between trees inferred with each given pair of models. Distributions are colored by median nRF and arranged in increasing order of median nRF. The vertical line through each distribution represents its median nRF, and rows showing `m1` comparisons are bolded for clarity. b) Proportion of PANDIT alignments whose the inferred tree, for each protein model, fell inside the `m1` confidence set, as assessed with AU tests. The percentage shown at the top of each bar indicates the percent of trees in the `m1` confidence set. c) Venn diagram depicting the number of PANDIT alignments whose trees inferred with `m4`, `m5`, and/or `JC` fell outside the respective `m1` confidence set of trees.

By contrast, nRF comparisons with `m5` and `JC` models were much higher, indicating that these two protein models tended to infer distinct topologies from `m1`–`m4` and `GTR` models. We did not observe a significant difference in mean nRF for the `m1`–`JC` and `m1`–`m5` comparisons, suggesting that `m5` and `JC` yielded trees with similar levels of deviation from `m1`. Interestingly, the mean nRF for the `m5`–`JC` comparison was significantly larger than were all other comparisons (*P* < 0.001). Therefore, while trees inferred with `JC` and `m5` models were fairly distant from `m1` trees, they were even farther from one another, indicating qualitative difference between `JC` and `m5` trees. Indeed, while both of these models fit the natural sequence data poorly, the poor fit of `JC` likely derived from their equal and therefore uninformative exchangeabilities, but the poor fit of `m5` models more likely derived from their misleading unequal exchangeabilities.

Further analysis with AU tests revealed that, for each alignment, most trees indeed fell in the `m1` confidence set of trees (Figure 6b). For the 200 alignments examined, 199 (99.5%) of `m2` trees, 194 (97%) of `m3` trees, and 198 (99%) of `GTR` trees fell in the `m1` confidence set of trees (*P* > 0.01). Therefore, although `m2`, `m3`, `GTR` models were poorer fits to the data compared to `m1`, trees inferred with these three protein models were statistically consistent with those inferred with `m1` models. By contrast, the `m4` models deviated from the `m1` confidence set somewhat more frequently, with only 183 (91.5%) of inferences consistent with the `m1` model. Finally, at least 1*/*3 of inferred trees under `m5` and `JC` each fell outside the `m1` confidence set of trees for their respective alignments.

We further asked whether the deviating trees inferred with `m4`, `m5`, and `JC` models represented the same or different PANDIT alignments, as depicted with the Venn Diagram in Figure 6c. There was relatively little overlap between which `m4` and `JC m5` trees fell outside the `m1` confidence set, but there was much more overlap between which `m5` and `JC` trees fell outside the `m1` confidence set. Even so, there were many instances where only the `JC` tree (from 19 alignments) or the `m5` tree (from 37 alignments) uniquely differed from the `m1` tree, again suggesting that `JC` and `m5` inferred qualitatively distinct phylogenies.

Unlike simulation results, where all `JC` trees fell inside the `m1` confidence set, many `JC` trees built from natural sequence data had significantly different topologies. We therefore suggest that further work is necessary to truly understand the performance of this simplistic yet potentially effective model. Indeed, based on Figure 6a, it appears that the equalrates `JC` model inferred unique topologies compared to any protein model with unequal exchangeabilities. While it is impossible to know which model(s), if any, converged upon the true phylogeny, the patterns observed from PANDIT data analysis imply that, so long as relative model fit is not exceptionally poor, the specific model is unlikely to strongly mislead or bias phylogenetic inference on protein data.

## Discussion

We have investigated whether relative model fit has a systematic effect on inference accuracy in single-gene protein phylogenetic inference. From both simulated and natural sequence data, we find that inferred topologies are highly robust to the fit of the employed protein model. These results were additionally robust to the simulation strategy, considering both generative models which violated and satisfied assumptions of empirical protein inference models. We emphasize that this study primarily considered the merits of relative model selection as a proxy for accuracy in phylogenetic topologies, but did not specifically consider accuracy in branch length estimation. As such, our results do not necessarily imply that relative model selection is wholly unimportant for phylogenetic inference. Instead, our results lead to the conclusion that relative model selection does not perform better than random chance at identifying which empirical protein model will yield the most reliable topology from amino-acid data.

Critically, the results presented here do not suggest that all protein models infer identical trees. The at-times wide distribution of nRF distances demonstrates that phylogenies inferred across varying models have, in many areas, distinct branching patterns (Figure 3, Figure 6a, and Figure S3). Instead, our results demonstrate that there is no clear, systematic shift towards more accurate inferences when relative model fit, as measured with information theoretic criteria, increases. While applying model selection procedures will not necessarily worsen a given analysis, there is similarly no robust evidence that applying relative model selection will improve analysis, as many users often presume. This is not to say relative model selection itself is either unnecessary or inaccurate, but rather that relative model selection is not a reliable “litmus test” for identifying which model will produce the most accurate and reliable inferences. In addition, because this study focused on relative model fit rather than absolute model fit, it remains a distinct possibility that all protein models had similar (potentially poor) absolute fits to the data. Future research endeavors should therefore assess the precise relationship between relative and absolute model fit measures to achieving a unified understanding of the merits of phylogenetic model selection approaches.

An unexpected but key finding in the simulation study presented here is that most models, regardless of relative fit, will recover similar proportions of highly-supported but incorrect nodes (Figure 4, Figures S3-9). This insight has important consequences for fundamental questions in phylogenetics and systematics. In particular, one reason it is desirable to avoid misspecified models is their presumed potential to yield supported but incorrect splits, or conversely correct splits which appear unsupported by the model (Sullivan and Joyce 2005). For example, recent studies aiming to disentangle fundamental relationships among mammals (Philippe et al. 2011; Moran et al. 2015; Tarver et al. 2016) and metazoans (Pisani et al. 2015; Ryan et al. 2013) have suggested that resolved phylogenies may have been elusive because many studies employed inadequate models which tend to yield strong yet inconsistent support. Our results imply that *all* protein models are likely to support incorrect splits, but no model is substantially over-represented for such splits.

Moreover, it has previously been observed that while the specific phylogenetic model used may not always impact topology, overly-simplistic models may have strong influences on measures of nodal support (Sullivan and Joyce 2005). We in fact did not observe this effect: The simplistic `JC` model tended to show similar levels of support compared to models with more complexity, considering MutSel and control simulations (Figure 4 and Figures S4-9). That said, the substantially larger differences in topologies inferred between `JC` and other models for empirical datasets obtained from PANDIT suggests any potential effect of model complexity on nodal support is simply not pronounced in simulated data. One potential reason for the comparable levels of nodal support between `JC` and more complex models in simulations is because, in fact, the models do not substantially differ in complexity because exchangeabilities are *a priori* fixed. Between any two commonly-used empirical protein models, the number of free parameters will usually differ by at most two: the proportion of invariant sites (`+I`) and/or a parameter representing ASRV such as the shape parameter of a discrete gamma distribution (`+G`). As such, the primary differences between any two empirical protein models are the specific values of their fixed substitution rates. This scenario is in stark contrast to nucleotide-level phylogenetic models, which are generally distinguished by the number of free parameters they consider. Indeed, the simplest nucleotide model contains only 1 free parameter, but the most complex models contain up to 12 free parameters, considering substitution rates along (Yang 2014). Therefore, while further work will improve our understanding of how protein models of varying complexity influence nodal support, it is possible that the relationship between model complexity and nodal support is mostly a concern for nucleotide-level data and associated nucleotide models.

In addition, we did not consider phylogenetic reconstruction from nucleotide data. However, a recent study by Abadi et al. (2019) used simulation to demonstrate that the `GTR+I+G` model and `JC` model as applied to nucleotide alignments do not produce systematically different phylogenetic topologies. Our results suggest that this phenomenon may also extend to protein-level data, ultimately providing increasing evidence that relative model selection is not predictive of accuracy in phylogenetic inference, in spite of decades of tradition. Similar to findings from Abadi et al. (2019), we also observed strong differences between the free-rate `GTR` and `JC` models as applied to natural sequence data, meaning that `JC` may not be as robust as simulations suggested.

Moreover, this study focused specifically on measures of topological accuracy when investigating how relative model fit might affect phylogenetic inference, and we did not investigate the effects of model fit on branch length inference. While such an analysis is clearly desirable, branch lengths under the simulation model (which operated at the codon level) and under inference models (which operated at the amino-acid level) are not directly comparable, and their relationship cannot be generalized. Under the MutSel simulation model, branch lengths are defined as the number of neutral codon changes per unit time, whereas under protein models used for inference, branch lengths are defined as the number of amino-acid changes per unit time, rendering these two quantities fundamentally distinct. Future efforts therefore may seek a unifying framework to compare branch lengths and divergence levels across model formulations.

Finally, this study focused exclusively on the practical ramifications of relative model selection when a single protein exchangeability models are applied to single-gene, non-partitioned data. We did not consider more complicated scenarios, such as the analysis of multiple concatenated genes in a partitioned analysis (Lanfear et al. 2017; Kainer and Lanfear 2015) and/or the use of more complex mixture models, such as the CAT model (Lartillot and Philippe 2004; Si Quang et al. 2008) or approaches that consider several exchangeability matrices proportioned across sites (Le et al. 2008; Huelsenbeck et al. 2008; Le et al. 2012). As mixture models have been shown to fit many datasets better than single exchangeability models, in particular for saturated or highly heterogeneous data (Le et al. 2008; Si Quang et al. 2008; Arenas 2015), future work should investigate whether the improvement in fit these complex models confer corresponds to qualitatively different phylogenetic inferences.

In sum, results presented here contribute to a growing body of evidence that the practical ramifications of model selection in phylogenetics may be vastly overstated. A key unanswered question in many of these findings is *why* there are such substantial differences in relative model fit even when these models perform extremely similarly. One possible explanation is that, while observed patterns in the data may more closely match some models than others, all available protein models may be similarly distant from describing the evolutionary process which in fact gave rise to the data. As such, while different models may better capture certain features of the data, none of them may have sufficient ability to capture the generative process of biological evolution. This insight may pave the way for the development of categorically novel modeling frameworks; if new protein exchangeability models are developed solely to improve the relative fit to data, but these new models do not yield meaningful consequences for inferences, the benefit to “building a better mouse trap” is modest at best.

## Acknowledgements

We thank Joseph Bielawski, Christopher T. Jones, Noor Youssef, Edward Susko, and Andrew J. Roger for helpful discussions about phylogenetic comparisons and uncertainty. We additionally thank the Associate Editor, one anonymous reviewer, and Shiran Abadi for valuable input to clarify findings in this study.

