## Supplementary Figures for "Relative model fit does not predict topological accuracy in single-gene protein phylogenetics"

Stephanie J. Spielman[1*]

[1]Department of Biological Sciences. Rowan University, Glassboro, NJ 08028.

#### Supplementary Table Descriptions

All tables are provided as separate CSV files.

**Table S1** All models `m1-m5` identified from simulated alignments.

**Table S2** Number of inferred trees with a Robinson-Foulds distance of 0 from the true simulation tree.

**Table S3** Number of inferred trees which differed significantly from the `m1` phylogeny by AU test.

**Table S4** All models `m1-m5` identified from PANDIT alignments.

### Supplementary Figures

##### Figure S1

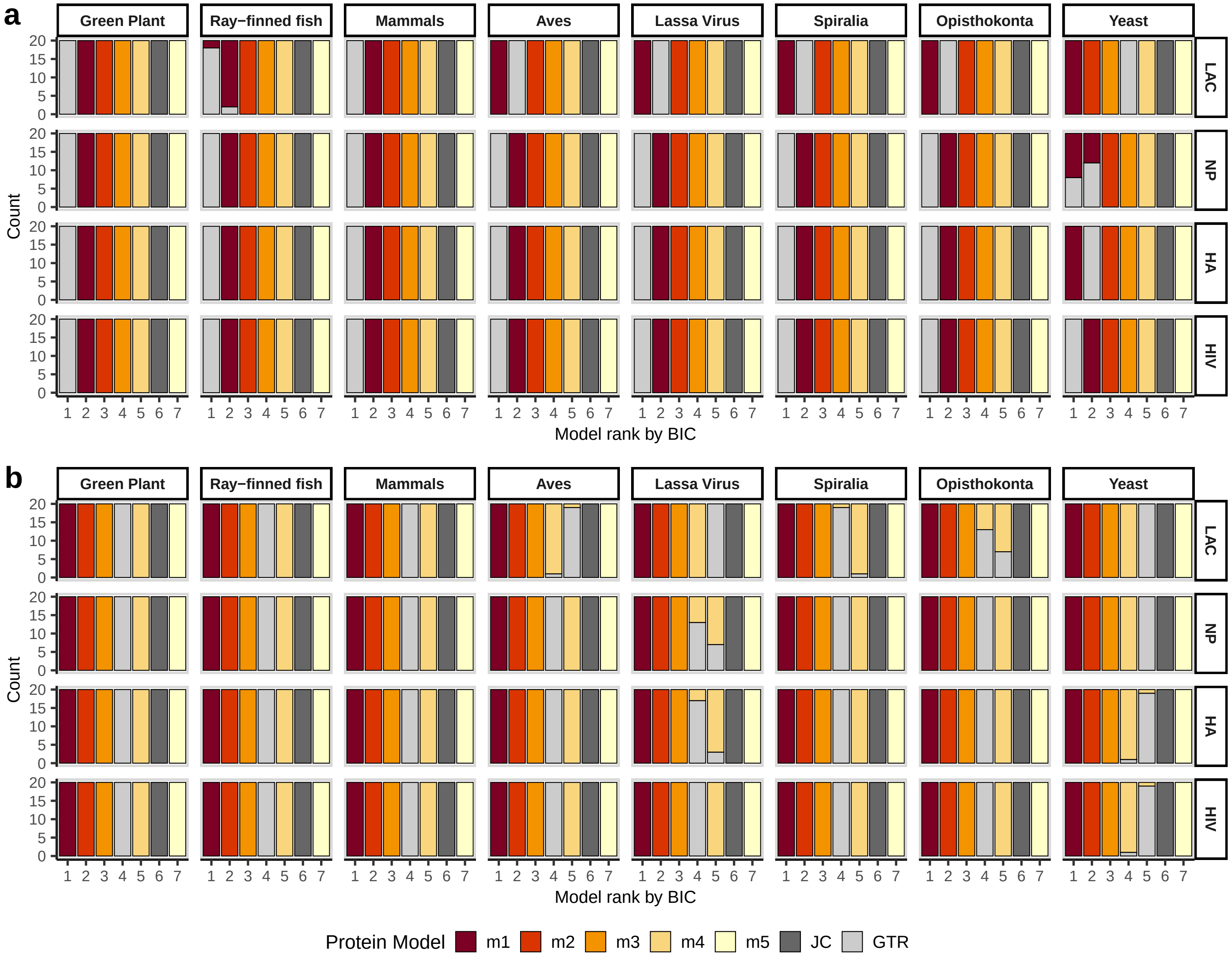


Relative rank of JC and GTR models, compared to the models `m1-m5`. The X-axis indicates the overall model rank among the seven models `m1-m5`, GTR, and JC. Each bar represents 20 simulation replicates, and each panel represents simulations for a given DMS parameterization and phylogeny. a) Results for for all MutSel simulations. b) Results for all control simulations.

#### Figure S2

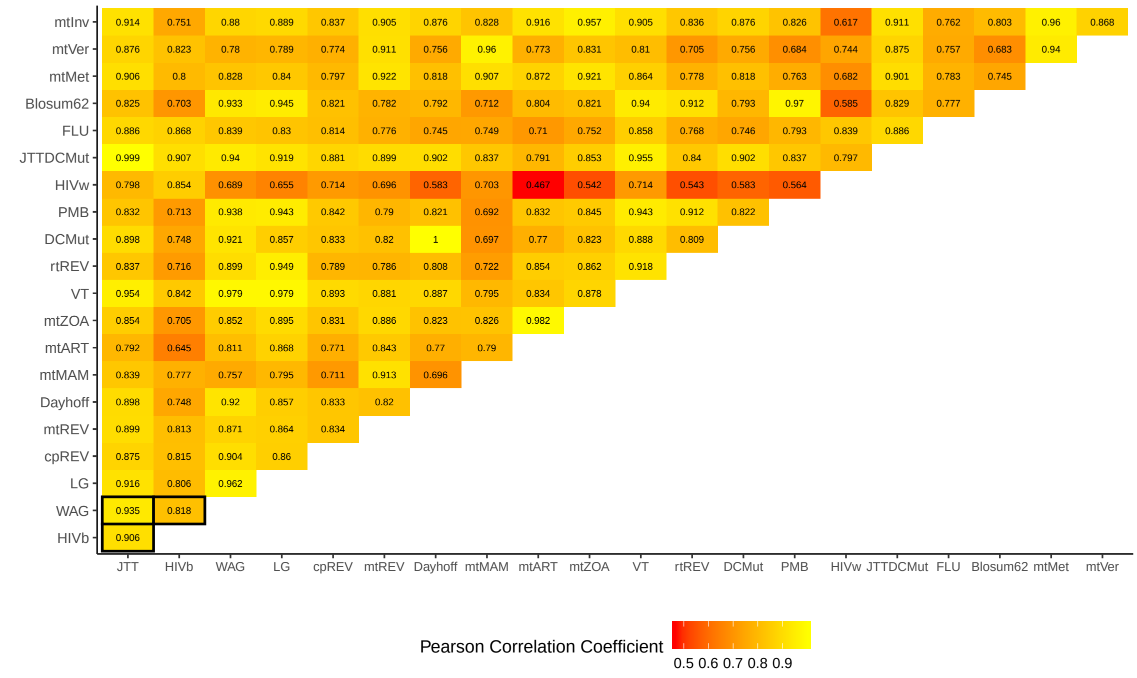


Pearson correlations among normalized exchangabilities for model matrices examined by `ModelFinder`. Comparisons between the MutSel simulation `m1` matrices recovered for simulations (JTT, HIVb, and WAG) are in boxes.

#### Figure S3

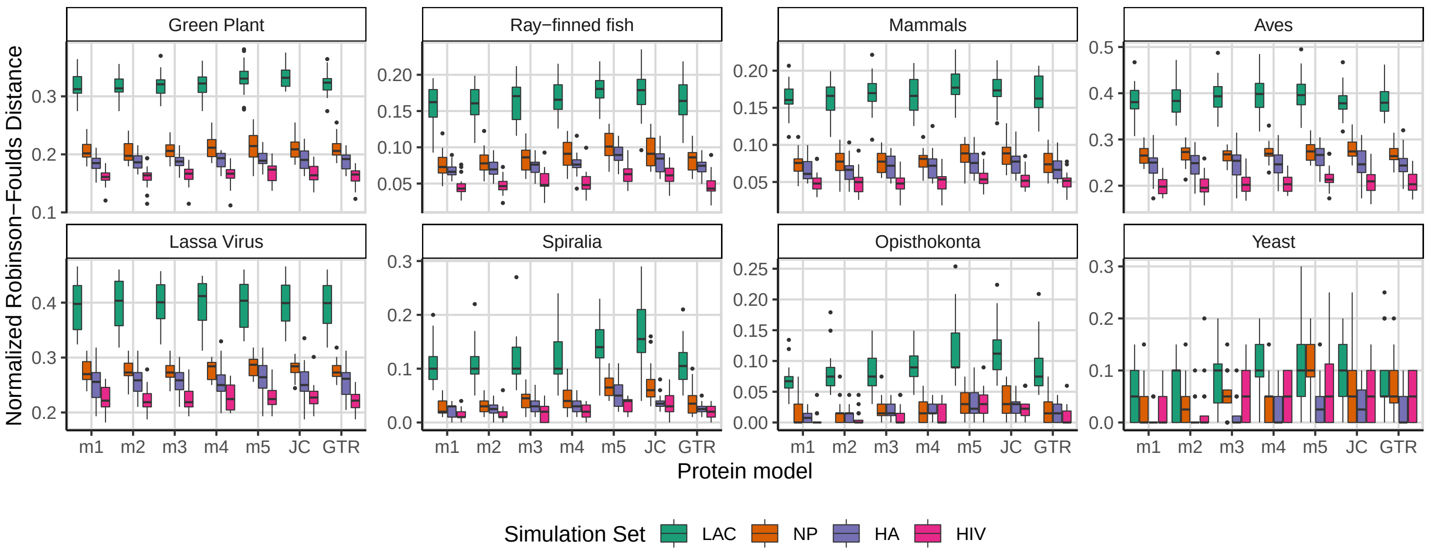


Normalized Robinson-Foulds distance (nRF) between tree inferences and the respective true tree from all control simulations. Each boxplot represents the distribution of nRF values for 20 simulation replicates.

**Figure S4**

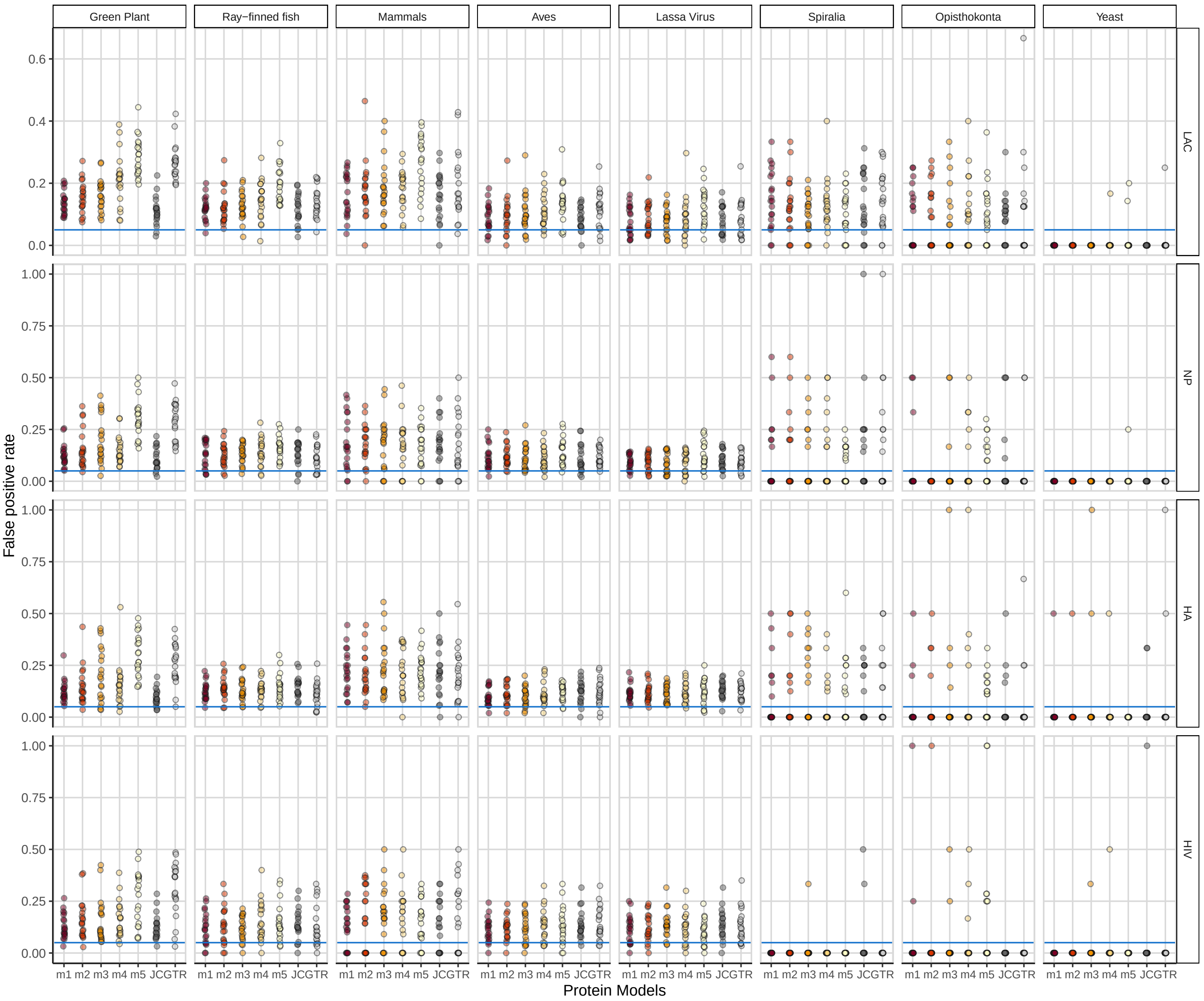


False positive rates (FPR) for node splits in tree inferences for all MutSel simulations, using 95% `UFBoot2` as a threshold. The horizontal line in each panel is the $y = 0.95$ line, representing the expected FPR.

#### Figure S5

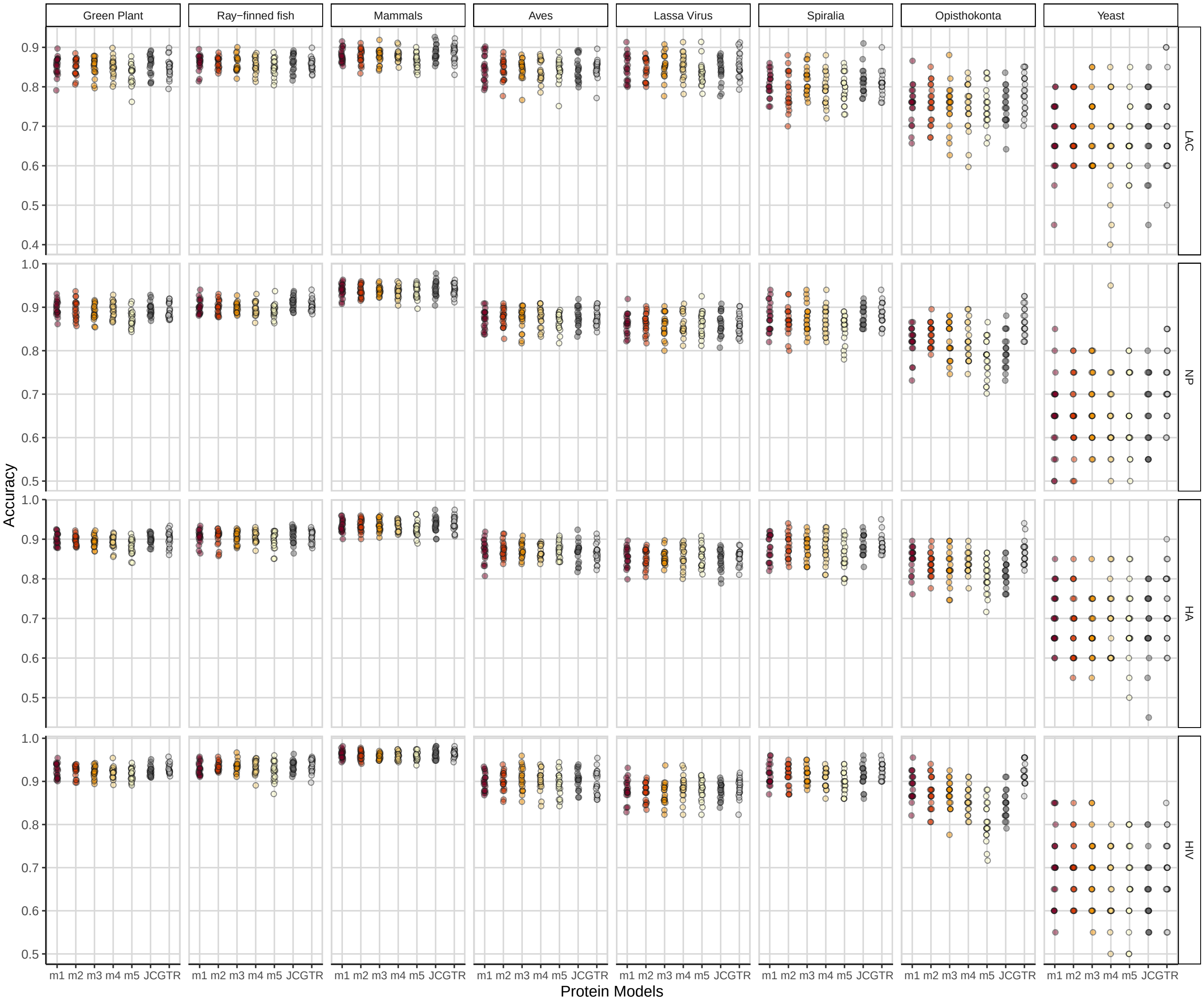


Proportion of accurately classified nodes in tree inferences for all MutSel simulations, using 95% `UFBoot2` as a threshold.

#### Figure S6

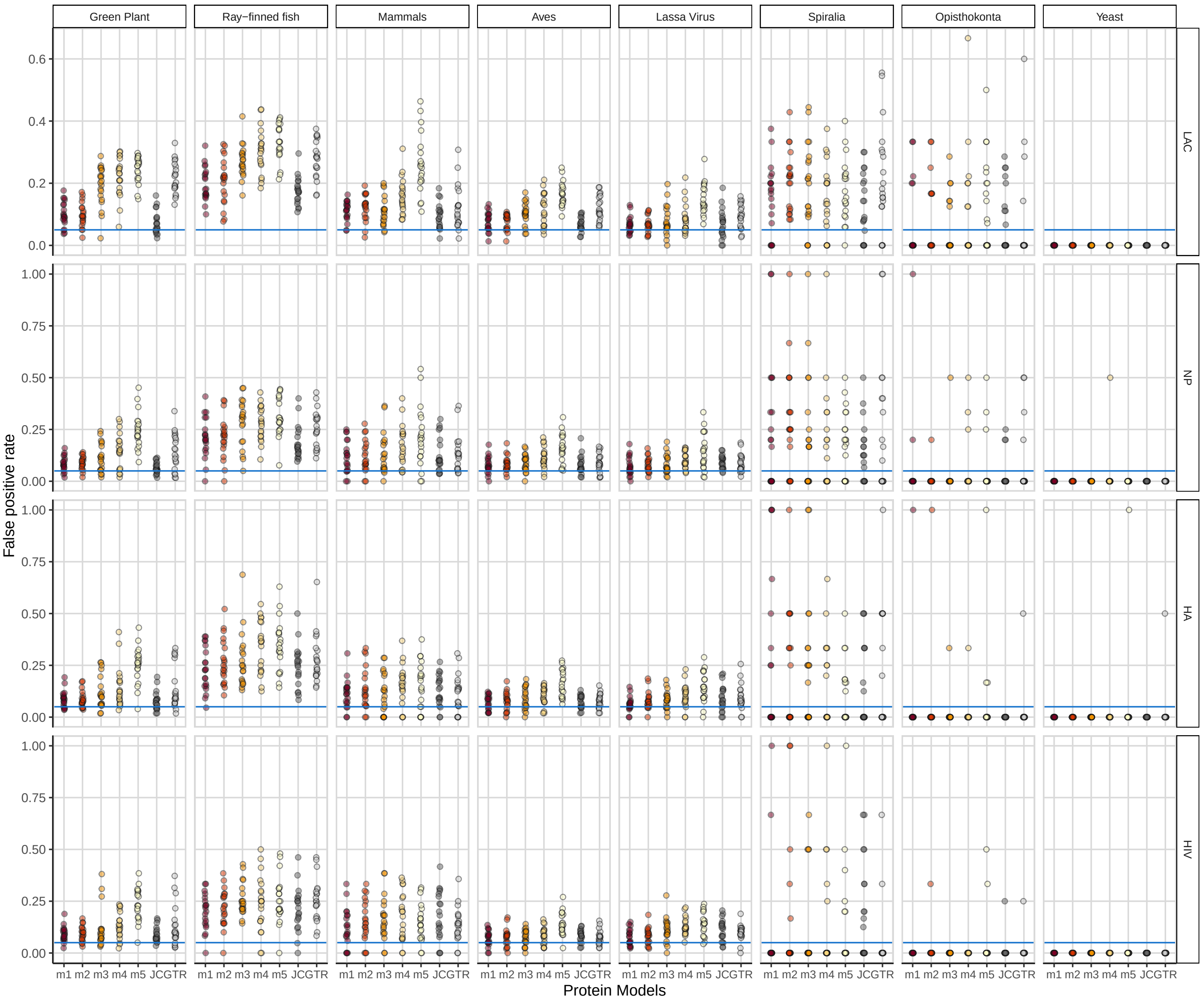


False positive rates (FPR) for node splits in tree inferences for all control simulations, using 95% `UFBoot2` as a threshold. The horizontal line in each panel is the $y = 0.95$ line, representing the expected FPR.

#### Figure S7

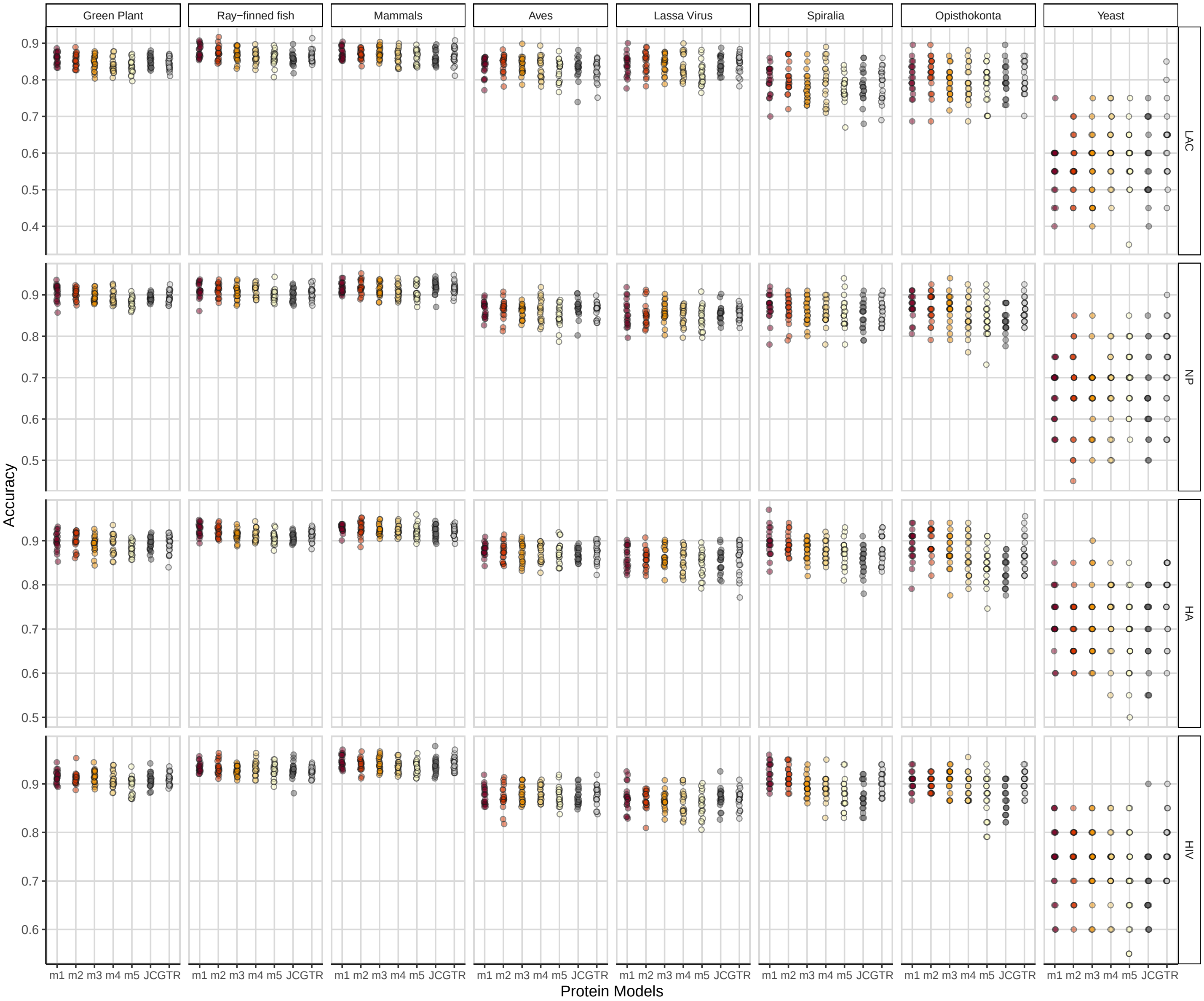


Proportion of accurately classified nodes in tree inferences for all control simulations, using 95% `UFBoot2` as a threshold.

#### Figure S8

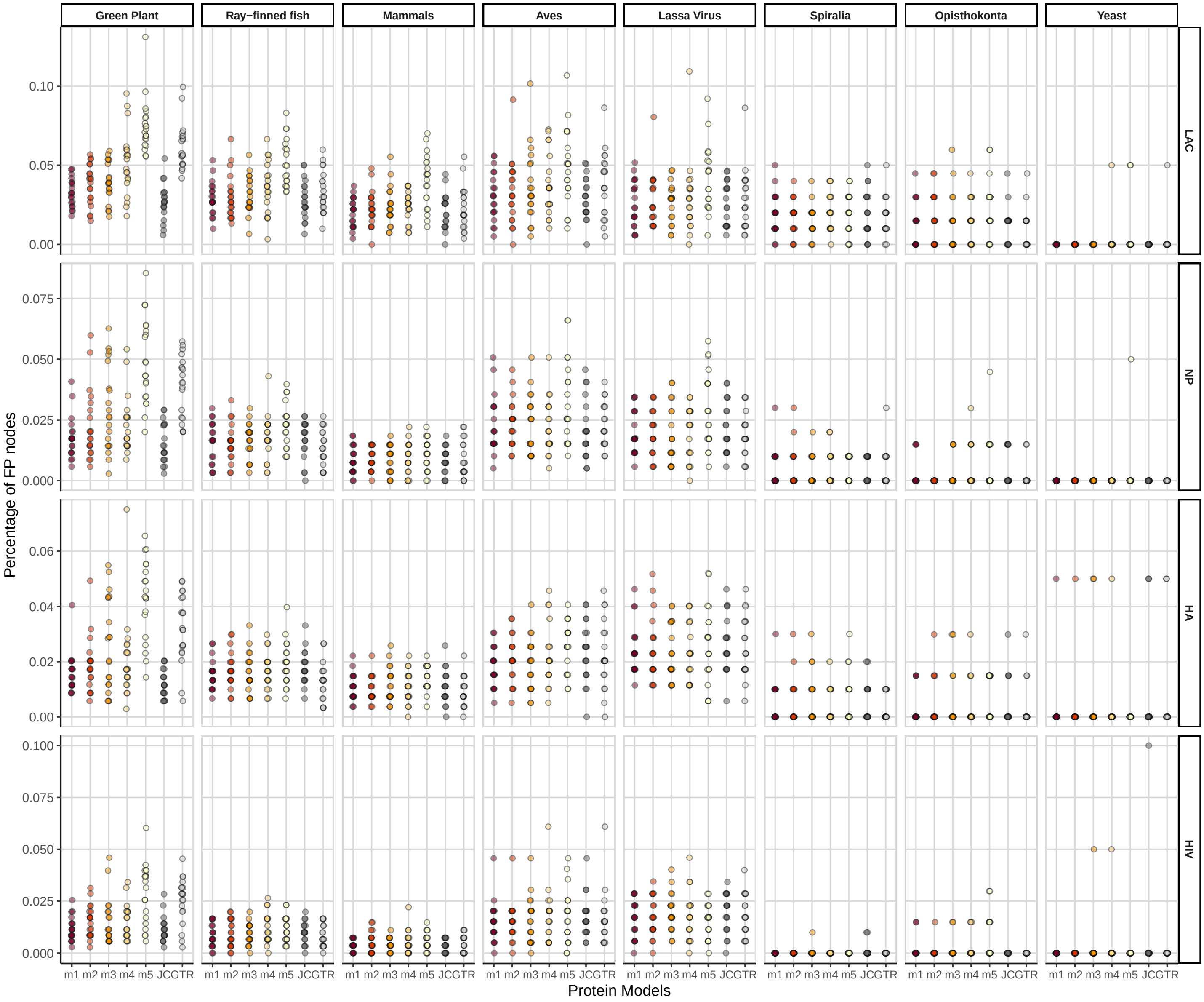


Proportion of all nodes in the tree that are false positives for all MutSel simulations, using 95% `UFBoot2` as a threshold.

**Figure S9**

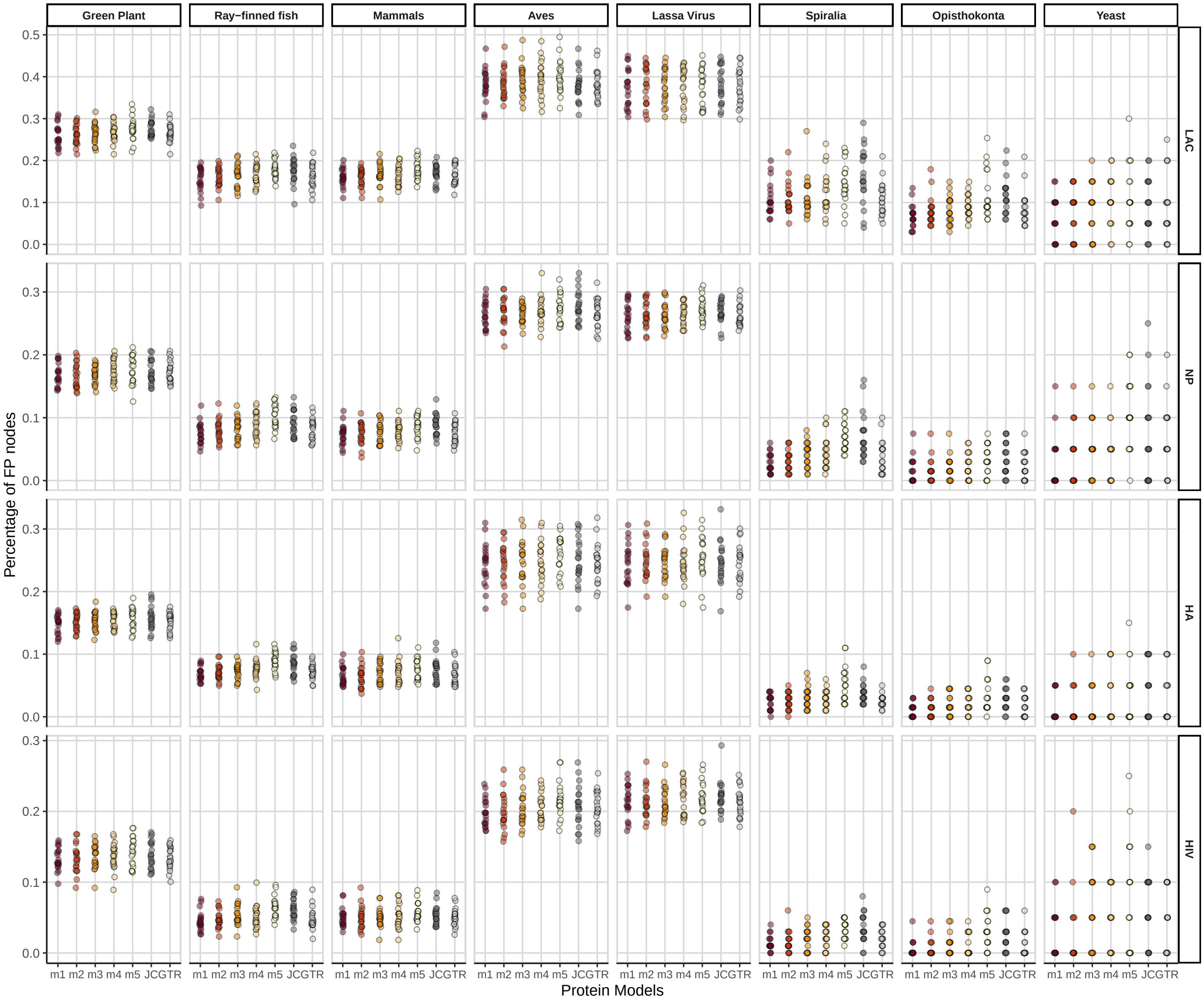


Proportion of all nodes in the tree that are false positives for all control simulations, using 95% `UFBoot2` as a threshold.
